# Immune determinants of pegivirus persistence, control, and cross-species infection in the laboratory mouse

**DOI:** 10.1101/2024.02.27.582314

**Authors:** Kylie Nennig, Teressa M. Shaw, Dave O’Connor, Jack Stapleton, Adam L. Bailey

## Abstract

Approximately 15% of the global human population is viremic with human pegivirus (HPgV), a +ssRNA virus in the *Flaviviridae* family. An unusual feature of HPgV is its ability to persistently infect individuals without causing overt disease or evoking robust immune responses, but this phenomenon is poorly understood due to a dearth of systems for studying HPgV. In this study, we create the first mouse model of PgV infection by adapting a PgV discovered in a wild rat (RPgV) to infect the standard laboratory mouse. Adaptation to the mouse initially required defective innate immunity and the accumulation of a single mutation in the E2 envelope glycoprotein, but passage into wild-type (WT) mice resulted in twelve additional mutations that enable persistent high-titer viremia, closely recapitulating the course of HPgV in humans. Mouse-adapted (ma)PgV infection of various knockout mice showed that lymphocytes exert a significant antiviral effect in the chronic phase of infection, but that this effect is also unable to fully control viremia in most individuals. Chronic type-I interferon signaling appears to paradoxically enable maPgV persistence, likely via T cell dysfunction that has been demonstrated in other chronic viral infections. However, unlike many persistent viruses, maPgV does not depend upon the induction of PD-1-mediated immune tolerance to maintain persistence. In-depth analysis of rare WT mice that achieved sterilizing maPgV immunity suggests that multiple possible paths to achieving PgV immunity exist and may include a combination of cellular, humoral, and non-canonical mechanisms. Altogether, our creation of maPgV opens up the vast murine toolkit for understanding the enigmatic biology of PgVs. In addition to novel insights into multiple aspects of PgV immunity, the lack of PD-1-mediated immune tolerance induced by PgV infection is unique among persistent viruses and suggests a highly novel mechanism of immune evasion.

**AUTHOR SUMMARY:** Viruses capable of persistently infecting an individual host have developed sophisticated mechanisms for evading host immunity, and understanding these mechanisms can reveal novel features of the host immune system. One such virus, human pegivirus (HPgV), infects ∼15% of the global human population, but little is known about its biology beyond the fact that it does not cause overt disease. We created the first mouse model of PgV infection by adapting a rat pegivirus to infect laboratory mice. This mouse-adapted virus (maPgV) caused infection that was detectable in the blood of mice for >300 days without causing signs of disease, closely recapitulating the course of HPgV in humans. This enabled unprecedented exploration of PgV immunity, revealing a pro-viral role for type-I interferon in chronic infection; a lack of PD-1-mediated tolerance to PgV infection; and multiple mechanisms by which PgV immunity can be achieved by an immunocompetent host. These data indicate that the PgV immune evasion strategy has aspects that are both common and unique among persistent viral infections. The creation of maPgV represents the first PgV infection model in wild-type mice, thus opening the entire toolkit of the mouse host to enable further investigation of persistent RNA infections.

## INTRODUCTION

The *Pegivirus* genus is one of four genera in the *Flaviviridae* family of enveloped +ssRNA viruses, which also includes the *Orthoflavivirus* genus (e.g., dengue, zika, yellow fever, West Nile), the *Hepacivirus* genus (e.g., hepatitis C virus), and the *Pestivirus* genus (e.g., bovine diarrhea virus) (*1*). Human pegivirus (HPgV, officially known as HPgV-1 and formerly known as GB virus C and also as Hepatitis G virus) causes long-lasting infection in ∼15% of the global human population, making it the most prevalent blood-borne RNA virus infection (∼10-fold more common than hepatitis C virus, HCV). HPgV is transmitted via sexual, blood-borne, and vertical routes, and causes a persistent viremia (as defined by the presence of ∼1×10^6-8^ viral genomes per mL of serum) that can last for decades, although ‘spontaneous’ clearance is well documented (*2*). Unlike other members of the *Flaviviridae*, HPgV does not cause overt disease. However, two noteworthy associations with HPgV infection have been identified: in large studies of HIV+ patients, HPgV co-infection is associated with a significant reduction in all-cause mortality and HIV-induced pathological immune activation, implying that HPgV infection is a positive prognostic indicator in the context of HIV infection (*3–6*); also, a weak but statistically-significant correlation exists between chronic HPgV infection and the development of non-Hodgkin B cell lymphoma (*7*).

Viruses capable of persistently infecting an immunocompetent host have sophisticated strategies for evading the host immune response. Common themes among viral persistence strategies include infection of immune-privileged sites (e.g., the central nervous system, CNS), latency, and the accumulation of mutations in key immune-targeting epitopes (i.e., immune escape) (*8, 9*). However, PgVs appear to employ none of these strategies, causing high-titer viremia for years without accumulating mutations indicative of immune escape (*10, 11*). Investigations into this apparently novel mechanism of immune avoidance have been hindered by the lack of systems for studying PgVs in the laboratory: no culture systems or small animal models of PgV infection exist.

In this study, we create the first small-animal model of PgV infection by adapting a PgV found in a wild rat (RPgV) to infect the standard laboratory mouse (c57BL/6). Adaptation to the mouse host initially required defective innate immunity (*STAT1*^-/-^ or *IFNAR*^-/-^), which resulted in the outgrowth of a minimally-mouse-adapted PgV containing just a single nonsynonymous mutation in the E2 glycoprotein. Upon passage into wild-type (WT) mice, this virus rapidly and consistently accumulated an additional 6 nonsynonymous mutations (concentrated in structural proteins) and 6 synonymous mutations (concentrated in non-structural proteins), resulting in a “fully” mouse-adapted PgV (maPgV). The kinetics of maPgV infection in WT mice was highly reproducible, with an initial peak of viremia at ∼15 days post infection (dpi) followed by the establishment of a viremic set-point around 100 dpi which lasted >300 days for ∼90% of infected animals. Infection of knockout mice lacking various immune genes showed that *RAG*^-/-^ mice maintained significantly higher levels of chronic-phase viremia, while *IFNAR*^-/-^ mice exhibited higher acute-phase viral loads but eventually cleared viremia at a significantly higher rate than WT mice. *PD1*^-/-^ mice displayed a blunted peak of maPgV viremia but established chronic viremic set-points similar to those of WT mice. Investigation of correlates of immunity in rare WT mice that controlled maPgV infection revealed distinct “pathways” to maPgV immunity, including cellular immunity that can be passively transferred between animals. Altogether, our novel creation of a mouse-adapted pegivirus greatly expands the questions that can be posed to interrogate the unique biology of this enigmatic group of viruses.

## RESULTS

### Disabling the type-I-interferon–Stat1 pathway enables adaptation of RPgV to mice

To leverage the vast resources available for studying mice, we set out to create a mouse model of PgV infection. Despite the prevalence of PgVs in mammals, naturally-occuring PgV infection of a house mouse (*Mus musculus*) has not, to our knowledge, been described. We therefore began by inoculating various immunocompromised mice––129 mice that lack Interferon (IFN)α and γ receptors (a.k.a “AG129”), *STAT1*^-/-^ c57BL6/J mice, and c57BL6/J mice expressing human STAT2 (*STAT2*^h/h^)––with pooled serum from HPgV-1-infected humans. None of these animals supported HPgV-1 infection, as determined by HPgV-1-specific RT-qPCR on serum (**Figure 1A**). PgVs appear to have a narrow host-species tropism: HPgV-1 will infect chimpanzees but not macaques (*10, 12*); however, baboon PgVs readily infect macaques (*11*). Given this, we inoculated wild-type (WT) and immunocompromised mouse strains with blood containing a PgV isolated from a closely related rat species. “Blips” of low-level viremia were detected in WT c57BL6/J and AG129 mice, but RPgV never achieved sustained titers of greater than 1×10^5^ genome copies (gc) per mL of serum in these animals (**Figure 1B**). In contrast, RPgV titers fluctuated erratically in three *STAT1^-/-^* mice until ∼60 days post-inoculation (dpi), after which RPgV titers rose to ∼1×10^9^ gc/mL of serum. This high titer was maintained until 88 dpi, at which point mice were euthanized and serum was pooled and diluted to create an infectious “mouse-adapted” PgV (maPgV) stock.

**Figure 1.**
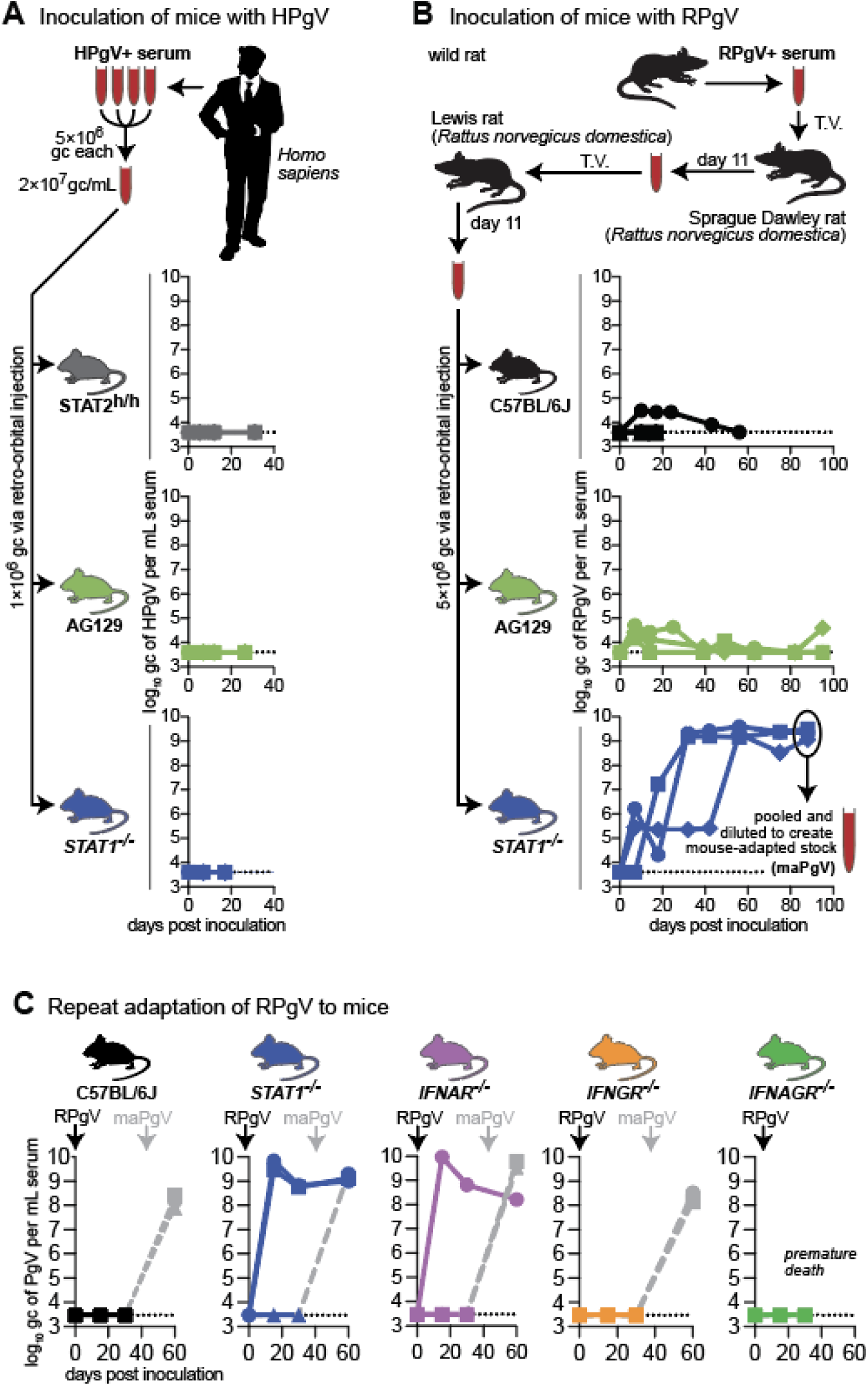
Disabling the interferon-alpha–Stat-1 pathway enables adaptation of RPgV to mice. **(A)** Pooled serum from multiple donors was inoculated into mice via retro-orbital injection: human STAT2-knock-in c57BL6/J mice (grey), *IFNAGR*^-/-^ “AG”129 mice (green), or *STAT1*^-/-^ mice (blue). Serum HPgV loads were measured using an HPgV-specific RT-qPCR assay, with the dashed line demarcating the limit of detection. **(B)** Serum containing RPgV was inoculated into mice via retro-orbital injection: wild-type c57BL6/J (black), *IFNAGR*^-/-^ “AG”129 mice (green), and *STAT1*^-/-^ mice (blue). Serum from 3 STAT1-knockout mice was collected and pooled to create a large “mouse-adapted” pegivirus (maPgV) virus stock. **(C)** Repeat of the RPgV adaptation study was performed via via retro-orbital injection of RPgV into: human wild-type c57BL6/J mice (black), *STAT1*^-/-^ mice (blue), *IFNAR*^-/-^ “AG”129 mice (purple), *IFNGR*^-/-^ “AG”129 mice (yellow), or *IFNAGR*^-/-^ “AG”129 mice (green). Re-inoculation with maPgV to examine RPgV-induced immunity is shown in gray.

Stat1 mediates signaling from both the IFNα/β receptor and the IFNγ receptor as well as several additional cell-type dependent receptors. To determine if the susceptibility of *STAT1^-/-^* mice to RPgV was mediated through a deficiency in IFNα/β or IFNγ signaling, we repeated the RPgV adaptation experiment in mice deficient in the IFNα/β receptor (*IFNAR*^-/-^), the IFNγ receptor (*IFNGR*^-/-^), and both types of IFN receptors (*IFNAGR*^-/-^) (**Figure 1C**). As expected, 0/3 WT mice supported RPgV infection, while 2/3 *STAT1*^-/-^ mice became infected with RPgV. 1/3 *IFNAR*^-/-^ mice became viremic with RPgV, while 0/3 *IFNGR*^-/-^ and 0/3 *IFNAGR*^-/-^ mice became infected, suggesting that PgV cross-species infection is restricted by type-I-IFN/Stat-1 signaling but is also a stochastic process. To determine whether the mice that remained RPgV-negative were immunologically naive, we inoculated these mice with our maPgV stock which resulted in robust infection in all animals, indicating that maPgV infection was unaffected by prior RPgV exposure.

### The kinetics of maPgV infection are highly reproducible and independent of route and dose

The route and dose of infection can have a profound impact on many aspects of viral infection including the kinetics of viral dissemination, disease, and the establishment of chronic infection versus the establishment of immune control. Thus, we undertook a dose × route study to examine whether our “mouse-adapted” PgV could cause persistent infection in WT mice, and what impact inoculating route and dose might have on this phenotype. We inoculated mice via retro-orbital (i.e., intravenous) injection with doses of maPgV ranging across 4 orders of magnitude (3×10^6^–3×10^2^ gc) (**Figure 2A**) or 2 orders of magnitude (3×10^6^–3×10^4^ gc) via intraperitoneal injection (**Figure 2B**) and followed viral loads in these animals for 300 days. All mice became infected, with viral loads detectable in all animals by 10 dpi. Viremia in intravenously-inoculated animals appeared to be more erratic and resulted in a few instances of viral clearance, which did not occur in intraperitoneally-inoculated animals. Although it appeared that animals inoculated with a lower intravenous dose had a higher propensity for eventually clearing their infection, repeat studies found this to not be the case (**Figure S1**). Ultimately, it appears that neither route nor dose has a significant impact on the trajectory of maPgV infection. Compiling data from all animals in the dose × route study shows that maPgV achieves peak viremia of ∼1.5×10^9^ gc/mL at ∼15-17 dpi, which gradually decreases to a set-point of ∼1×10^6^ gc/mL by ∼100 dpi. Approximately 10% of animals control their viremia to undetectable levels during this time, with most instances of control occurring between 100–150 dpi.

**Figure 2.**
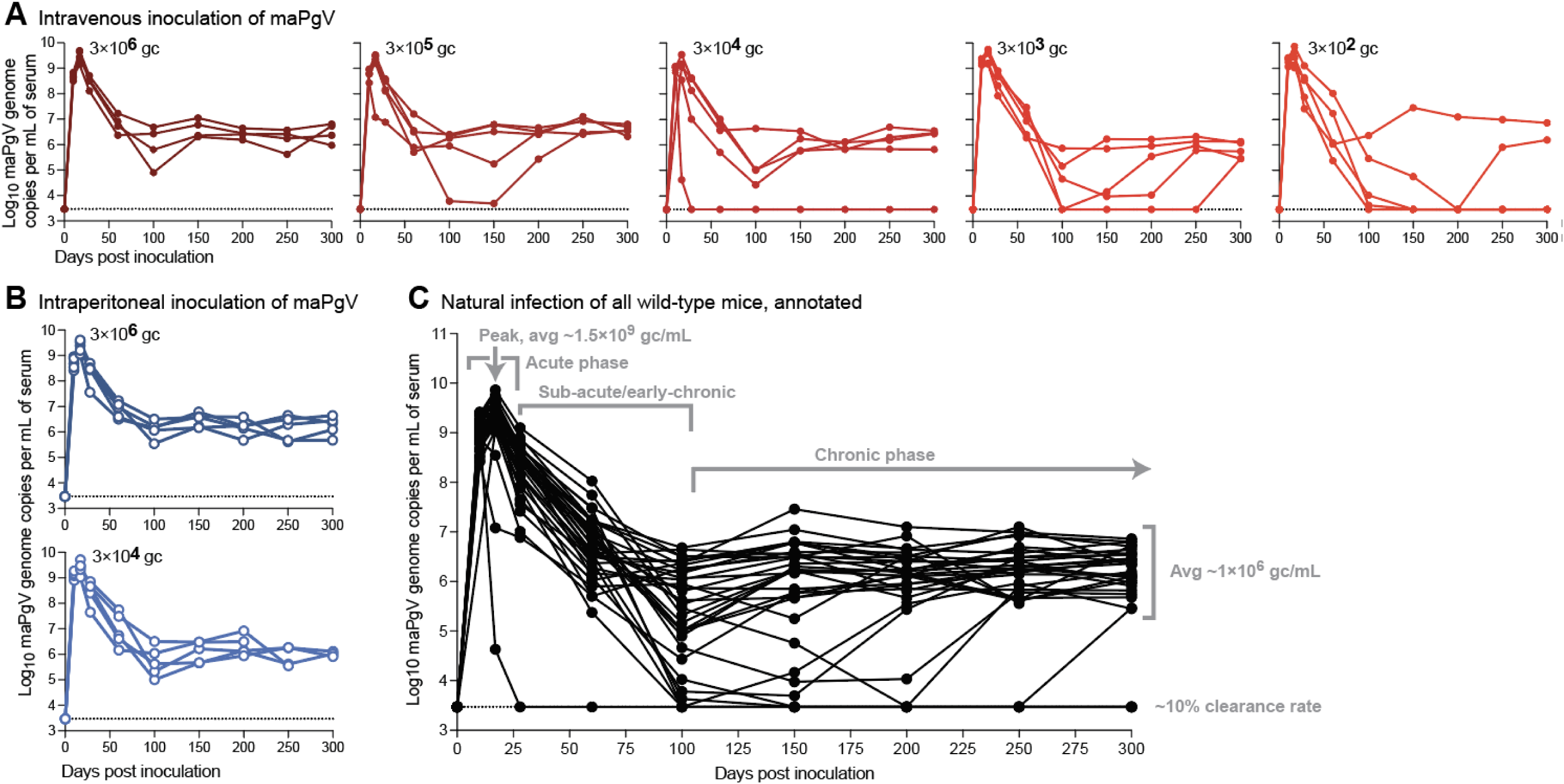
Persistent infection of wild-type mice with maPgV. Serum viral loads in wild-type mice inoculated with maPgV via (**A**) retro-orbital (i.e., intravenous) injection (red) or (**B**) intraperitoneal injection (blue). (**C**) Viral loads of all animals in the route × dose study, with key phases of infection annotated in gray. Limit of detection for the viral load assay is shown as a dashed line.

### A single mutation in the E2 envelope glycoprotein (R80L) is important for murine adaptation of RPgV

To examine the adaptive mutations that enable productive RPgV infection of mice, we performed an initial “unbiased” deep sequencing characterization of our maPgV stock. The incomplete *de novo*-assembled maPgV genome was then used as a launching point for performing 5′ and 3′ rapid amplification of cDNA ends (RACE) to generate the complete end-to-end maPgV genome sequence. Using the complete maPgV genome sequence, we then used pooled primer sets to generate overlapping amplicons that yielded deep sequencing coverage of >99% of the genome for most samples. Deep sequencing of the original RPgV serum sample revealed very little intra-host diversity, with none of the subsequent mutations identified in mouse-adapted PgV descendants observable at >5% frequency (**Figure 3**). Upon inoculation of this nearly-clonal virus into *STAT1*^-/-^ or *IFNAR*^-/-^ mice, mutations accumulated throughout the genome. Non-synonymous mutations clustered primarily in structural genes, predominantly E2 and X, although viruses in one *STAT1*^-/-^ and one *IFNAR*^-/-^ mouse also accumulated non-synonymous mutations in NS5A and NS5B. Synonymous mutations accumulated throughout the non-structural genes, and there were few mutations that arose in the 5′ or 3′ UTR. Of the multiple murine-adapted descendant viruses, several shared common mutations. However, only one mutation––a non-synonymous mutation in the putative E2 glycoprotein (R80L)––was common to all minimally-murine-adapted PgVs. Of note, only the pooled “maPgV stock” caused robust infection in WT mice; the other minimally-murine adapted PgV descendants caused low-titer viremia in WT mice that was below the threshold required for sequencing.

**Figure 3.**
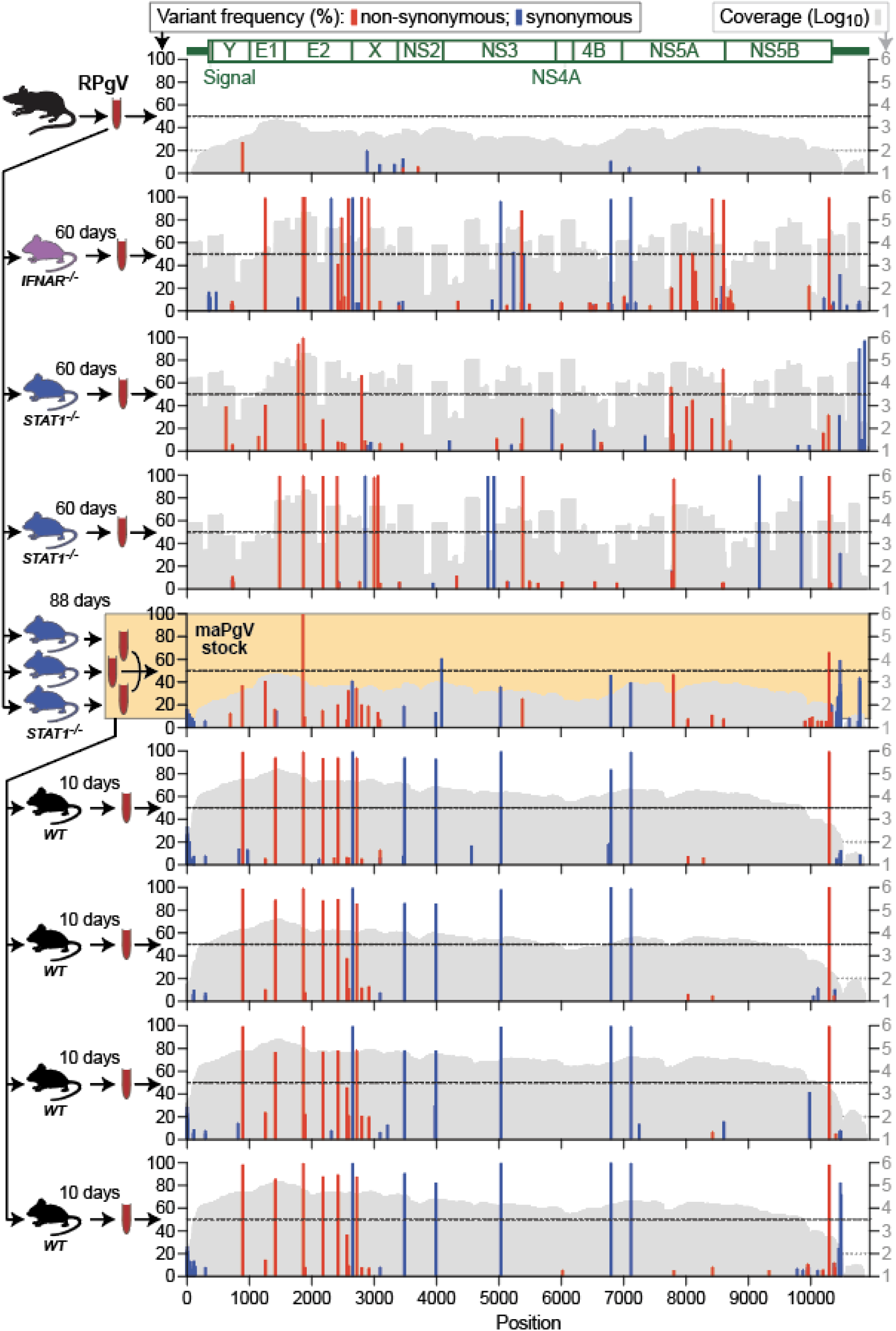
A single mutation in the E2 envelope glycoprotein (R80L) is important for murine adaptation of RPgV. Illumina deep sequencing of RPgV at various points during mouse adaptation. The genome position of RPgV/maPgV is shown along the X-axis, with a schematic of predicted mature proteins shown in green across the top. The frequency of non-synonymous mutations (red) and synonymous variants >5% relative to the RPgV consensus sequence are shown along the left Y-axis, with a dashed black line denoting 50% frequency (i.e., consensus-level variants). Coverage is shown in gray on a log10 scale along the right Y-axis with a read-depth cutoff of 100 shown as a gray dashed line, below which variants were not called. The pooled “maPgV stock” described in Figure 1 is highlighted in yellow. Note that some samples were sequenced via unbiased deep sequencing and other were sequenced by primal sequencing, generating the “mountainous” versus “city-scape” appearing coverage plots, respectively.

### Thirteen mutations confer full murine adaptation of RPgV

Passage of the pooled maPgV stock into WT mice resulted in the rapid and consistent accumulation of 12 additional mutations, with non-synonymous mutations clustered in structural genes (except for one mutation in NS5B, the putative RNA-dependent RNA-polymerase) and synonymous mutations in non-structural genes (except for one silent mutation in E2) (**Figure 4** and **Figure S2**). The accumulation of these additional 12 mutations in an immunocompetent host indicated that they might be important for evading additional aspects of the immune system. To test this hypothesis, we infected *IFNAR^-/-^*and *RAG*^-/-^ mice (which lack mature T and B cells) with the maPgV stock and sequenced virus at peak viremia (15 dpi for *IFNAR^-/-^* and 200 dpi for *RAG*^-/-^). Remarkably, in both groups of mice, all 13 of the fully mouse-adapting mutations rose to consensus levels without the consistent accumulation of additional mutations. Although the functional implications of these mutations remain to be determined, it seems clear that this set of 13 mutations confers full adaptation to the murine host and are likely selected for their enhanced ability to interact with murine host factors.

**Figure 4.**
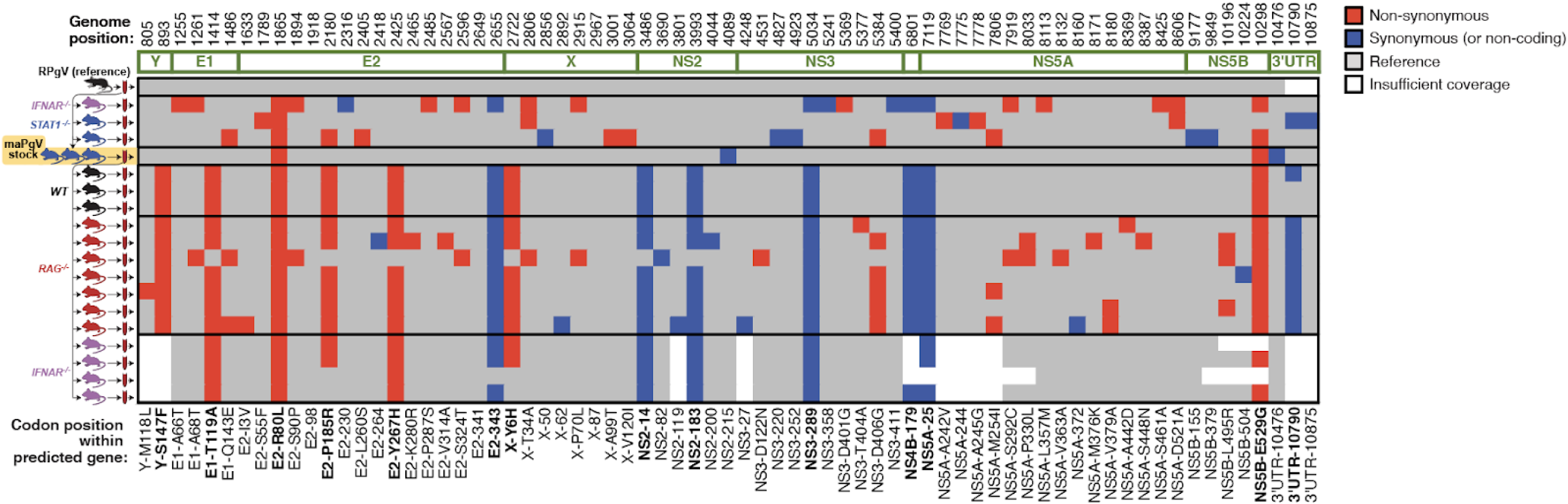
Thirteen mutations confer full murine adaptation of RPgV. Summary of all consensus-level variants detected in the expanded sequencing dataset. Equivalent analysis to that shown for a subset in Figure 3 can be found in Figure S2. Non-synonymous and synonymous variants are shown in red and blue, respectively. Coverage > 100 reads is shown in gray. The thirteen mutations that consistently accumulate during adaptation to the murine host are bolded along the bottom.

### Type-I interferon signaling contributes to PgV persistence

PgV persistence remains a poorly understood phenomenon, without a clear understanding of how various components of the immune system impact the level and duration of PgV viremia. To investigate this, we first examined the role of type-I interferon (IFN)––a master regulator of antiviral immunity––on maPgV infection by infecting mice deficient in the IFNα receptor (*IFNAR*^-/-^). Compared to maPgV viremia in WT mice, *IFNAR*^-/-^ mice displayed a similar peak of viremia but maintained higher viral loads in the sub-acute/early-chronic phase of infection (**Figure 5A,B**). Interestingly however, a majority of *IFNAR*^-/-^ mice went on to clear maPgV infection by 200 dpi, compared to a clearance rate in WT mice of only ∼10%, suggesting that intact type-I IFN signaling in late-chronic PgV infection facilitated PgV persistence. This is consistent with prior research on chronic viral infections, which also found that IFNAR signaling blockade restored T cell function (*13–15*). IFNγ can also have a significant impact on chronic viral infection, and so we examined the effect of IFNγ on maPgV infection by infecting mice deficient in the IFNγ receptor (*IFNGR*^-/-^), along with mice deficient in both IFNα and γ receptors (*IFNAGR*^-/-^). In contrast to *IFNAR*^-/-^ mice, *IFNGR*^-/-^ mice displayed a blunted acute-phase maPgV peak followed by a slightly elevated chronic-phase set point without any instances of viral clearance. *IFNAGR*^-/-^ mice had an acute-phase maPgV trajectory similar to *IFNAR*^-/-^ mice, but animals diverged into one of two patterns in chronic phase, either sustaining very high titers or progressing towards viral clearance. We also examined the impact of adaptive immunity on maPgV infection by infecting mice deficient in T and B cells (*RAG*^-/-^), which consistently sustained high levels of maPgV viremia.

**Figure 5.**
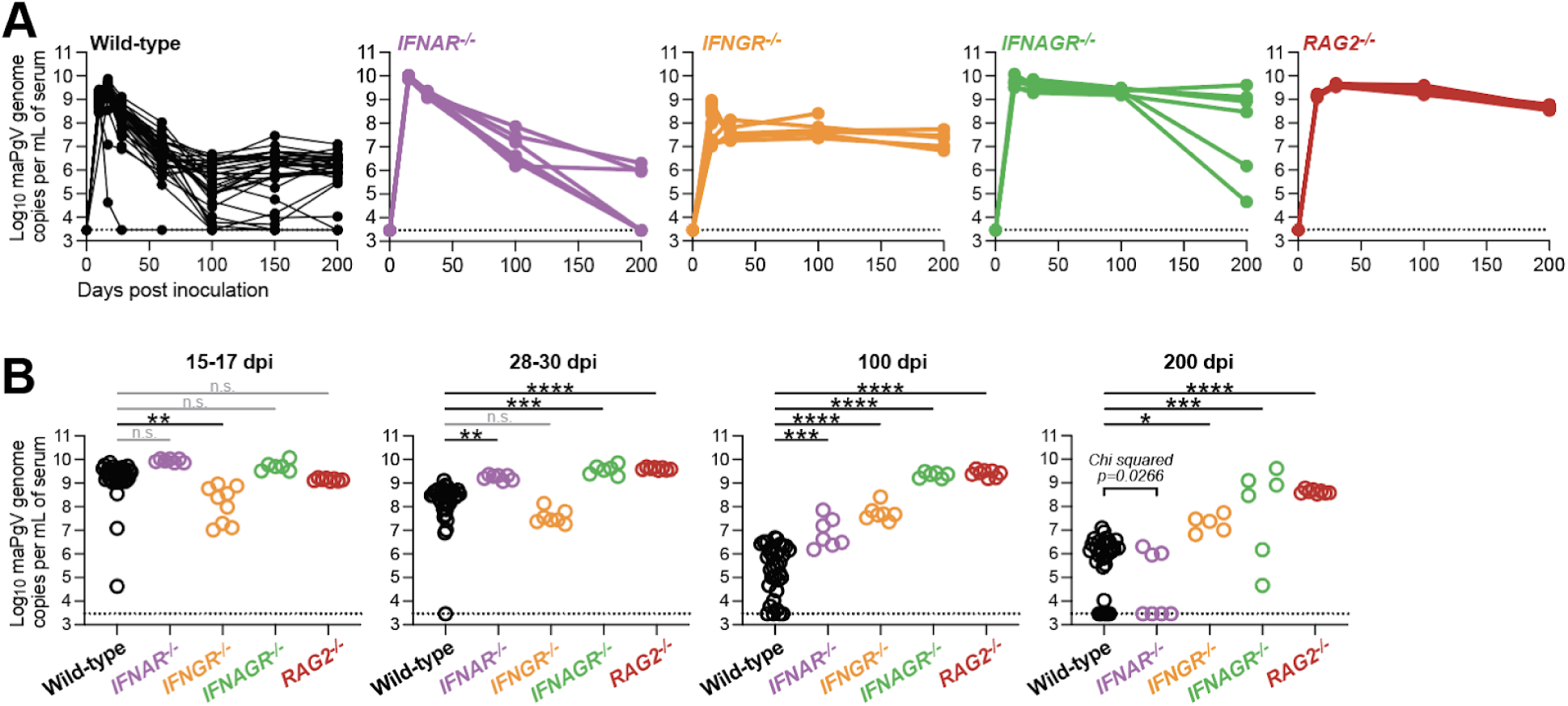
Type-I interferon signaling contributes to PgV persistence. (**A**) Serum viral loads of maPgV in various immunocompromised mouse strains over time (wild-type: black; *IFNAR*^-/-^: purple; *IFNGR*^-/-^: orange; *IFNAGR*^-/-^: green; *RAG*^-/-^: red). (**B**) Comparison of maPgV serum viral loads in immunocompromised mice strains at various time points during infection using one-way ANOVA in relation to wild-type mice, corrected for multiple comparisons (*:p≤0.05; **:p≤0.01; ***:p≤0.001; ****:p≤0.0001). Note: data from Wild-type mice is from the same data-set shown in Figure 2; data from immunocompromised mice is combined from two independent cohorts.

### PgVs do not require PD-1-mediated immune tolerance for persistence

The induction of immune tolerance plays a well-described role in the establishment and maintenance of persistence in several viral infections (*16*). In particular, upregulation of programmed cell death protein-1 (PD-1), which inhibits the activation of virus-specific T lymphocytes (among other, less well-described functions), plays a central role in the induction of tolerance, blunting the anti-viral immune response to prevent immunopathology at the cost of persistence (*17, 18*). Indeed, mice deficient in the PD-1/PD-L1 signaling axis succumb to fatal immunopathology when infected with lymphocytic choriomeningitis virus (LCMV, clone 13), a commonly used model of RNA virus persistence (*19*). To determine if induction of PD-1 mediates PgV persistence––without any way of measuring PgV-specific T cells––we infected *PD1^-/-^* mice with maPgV. *PD1^-/-^*mice exhibited slightly lower acute-phase (15 dpi) viral loads compared to WT mice, but this difference disappeared as infections progressed (**Figure 6A**). *PD1^-/-^* maPgV-infected mice also displayed no signs of disease or impaired weight gain relative to age/sex-matched naive *PD1^-/-^* mice, naive WT mice, or maPgV-infected WT mice (**Figure 6B**). In contrast, *PD1^-/-^* mice infected with LCMV (clone 13) rapidly lost weight, as described previously.

**Figure 6.**
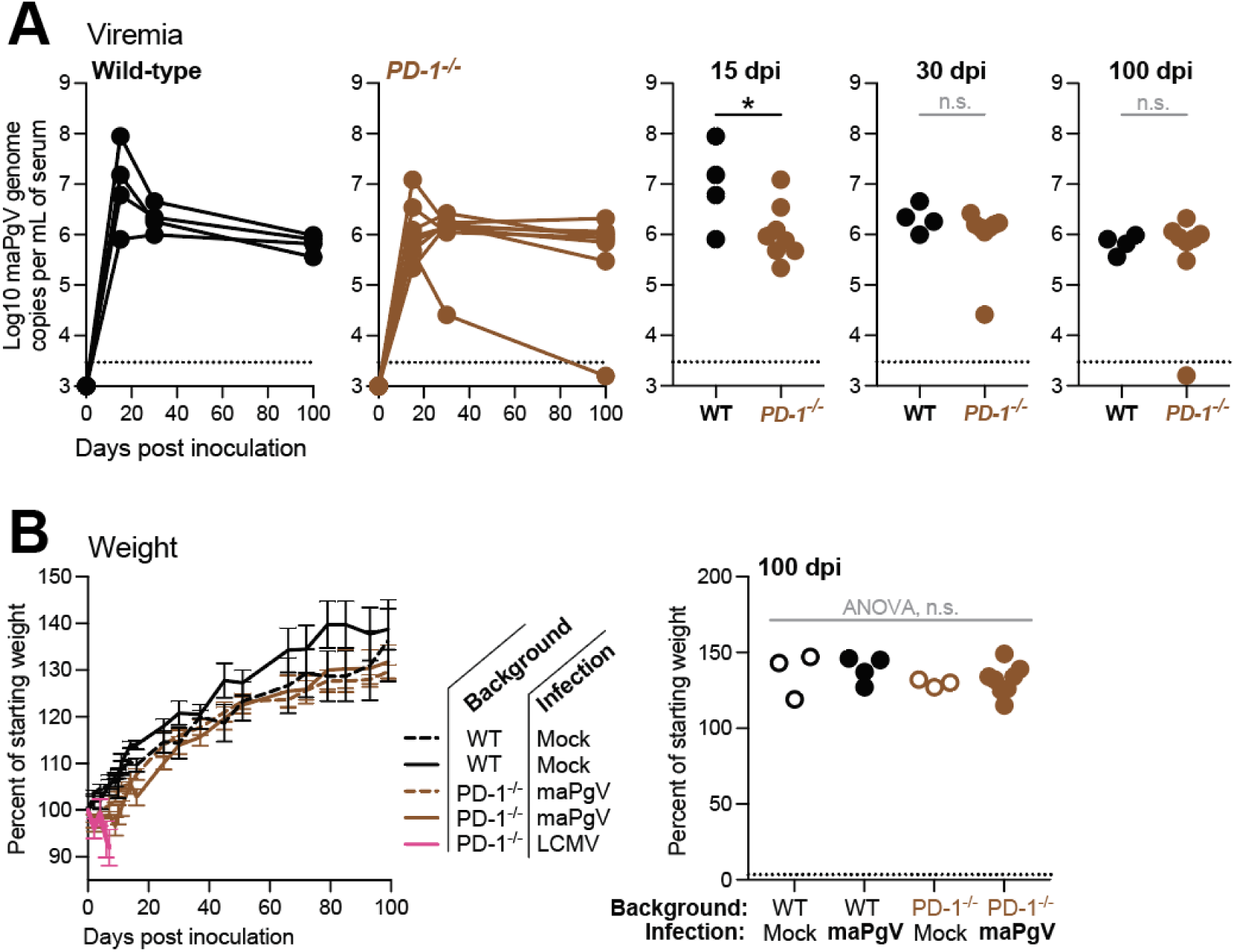
PgVs do not require PD-1-mediated immune tolerance for persistence. (**A**) Serum viral loads of maPgV in wild-type (black) versus *PD-1*^-/-^: (brown) with unpaired t-test used to determine significance (*:p≤0.05). (**B**) Weight gain/loss data for wild-type (black) and *PD-1*^-/-^ (brown) infected with mock (dashed line or open circle), maPgV (solid line or solid circle), or LCMV (pink).

### Natural PgV immunity can be achieved via multiple immune mechanisms

Approximately 10% of mice infected with maPgV control viremia to undetectable levels by 200 dpi. Most of these animals achieve undetectable viremia between 100-150 dpi, but we also observed one example of rapid maPgV clearance immediately following a typical acute-phase peak. To examine correlates of immunity in these relatively rare “controllers,” we first re-challenged these animals with maPgV, which leads to “blips” of viremia in 2/4 animals that were again driven to undetectable levels (**Figure 7A**). We then harvested splenocytes and serum from these controllers and transferred these tissues into PgV-naive mice. Two recipient mice were used for each donor×tissue, and each recipient mouse had been sublethally-irradiated 4 days prior to enable engraftment of adoptively-transferred cells. Two days after tissue transfer recipient mice were inoculated with maPgV and peak viremia was assessed at 15 dpi. Results varied by donor and tissue. Tissues from one maPgV-immune donor had no impact on acute-phase maPgV load; serum and splenocytes from a second donor both exhibited a modest anti-maPgV activity; while splenocytes from a third donor showed near-sterilizing maPgV immunity (**Figure 7B**). To further test the limits of the anti-PgV immunity mediated by splenocytes from this unique donor, we adoptively transferred cryopreserved splenocytes from this donor into two Rag^-/-^ mice that had been infected for >250 days. Within 15 days post-transfer, maPgV viral loads had decreased by three orders of magnitude, from a set-point of ∼1×10^8-9^ gc/mL to ∼1×10^5-6^ gc/mL (**Figure 7C**). In contrast, transfer of splenocytes from a maPgV-naive donor had no impact on chronic phase viral loads. By 20 days post-transfer, maPgV viremia was undetectable in Rag^-/-^ mice that had received anti-maPgV splenocytes. These mice remained aviremic, even after maPgV rechallenge.

**Figure 7.**
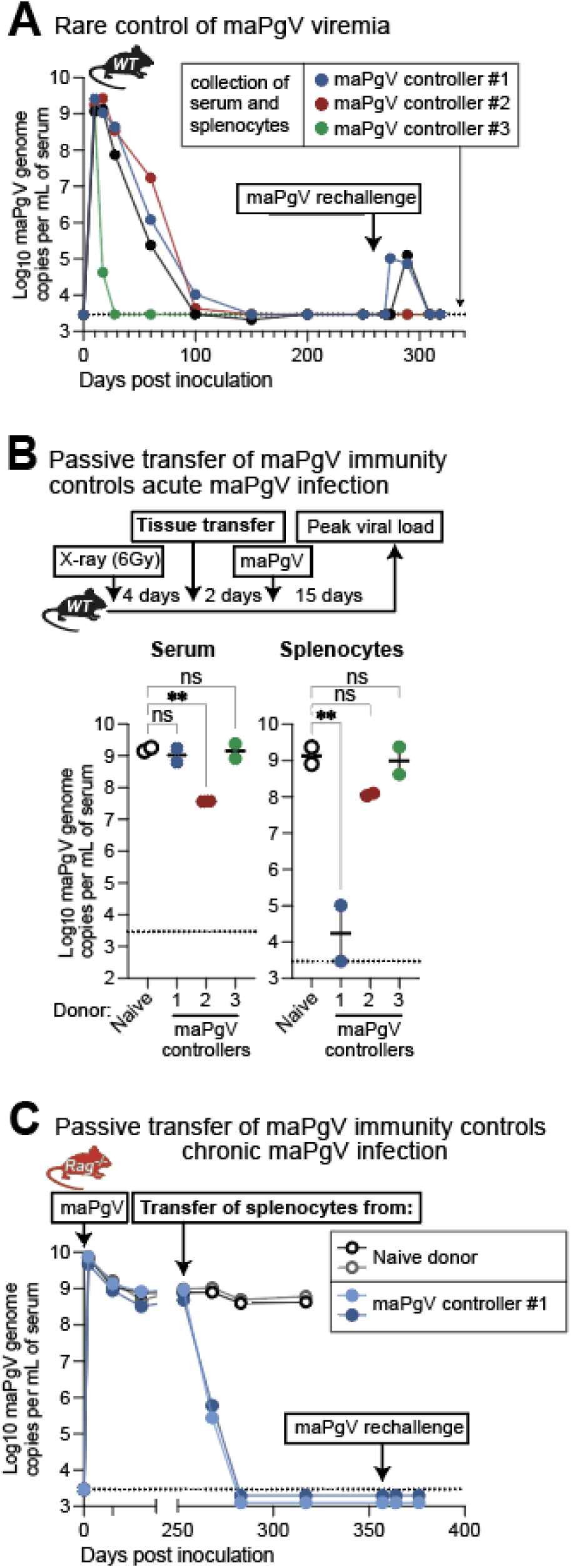
Natural PgV immunity can be achieved via multiple immune mechanisms. (**A**) Trajectories of viremia in 4 mice who ultimately cleared maPgV infection. (**B**) Serum and splenocytes from 3 animals that cleared maPgV viremia were collected and adoptively transferred into naive wild-type mice that had been mildly immunocompromised via X-ray irradiation (two recipients per donor; colors correspond to donors in part A). At the typical peak of maPgV viremia (15 dpi), viral loads were measured and compared to controls that received serum or splenocytes from a maPgV-naive wild-type mouse using one-way ANOVA with correction for multiple comparisons (**:p≤0.01). (**C**) Cryopreserved splenocytes from one donor (blue, part A) were adoptively transferred into two chronically-infected *RAG*^-/-^ mice (closed circles), with transfer of splenocytes from a naive donor (open circles) serving as controls.

## DISCUSSION

The laboratory mouse has become the dominant animal model for studies in virology and immunology for a multitude of reasons (cost, size, housing requirements, reproductive time, investments in transgenic and reagent technologies, ethical considerations, etc). However, there are relatively few mouse models of chronic viral infection and many of these viruses require an immunocompromised host for persistent viral replication. Additionally, no immunocompetent mouse models exist for studying infections by viruses in the related *pegi*-, *hepaci*-, or *pesti-virus* genera (*20*). In this context, our creation of a mouse-adapted pegivirus (maPgV) represents a significant advance, as maPgV causes highly-reproducible persistent infection in wild-type mice, opening up the entire toolkit of the murine host for further exploration of maPgV as a *bona fide* model of chronic infection. The close relatedness of maPgV to human PgV (HPgV)––both in terms of phylogenetic ancestry and their similar course of infection in mice and humans, respectively––is another noteworthy aspect of the maPgV model given that many viruses used as chronic infection models in mice do not have a directly-relevant human counterpart.

Our infection of mice with RPgV, but not HPgV, adds to a growing body of evidence indicating that PgVs have a narrowly-restricted species-tropism (*10, 11*). Although the molecular interactions governing PgV species-tropism remain unknown, the unique susceptibility of innate-immune-compromised (*STAT1*^-/-^ and *IFNAR*^-/-^) mice to RPgV suggests that innate immune effectors act to restrict cross-species PgV infection. Thus, we propose that RPgV adaptation to the murine host occurred through the following mechanism: first, in the absence of innate immune restriction, RPgV was able to establish inefficient/low-level infection in mice. This then allowed RPgV to acquire additional mutations that were selected for enhanced/efficient interactions with the murine host, many of which reside in the envelope glycoproteins. Although attributing specific mutations found in the fully-mouse-adapted (ma)PgV to specific host interactions will be a major undertaking, future comparisons should be able to draw upon the RPgV and maPgV genome sequences to explore interactions between these viruses and their respective hosts.

The immune response to PgVs has often been conceptualized as weak or even non-existent, as evidenced by the ability of PgVs to persistently infect immunocompetent hosts without accumulating mutations indicative of adaptive immune pressure/escape (*11, 21, 22*). Nevertheless, *RAG*^-/-^ mice sustain chronic-phase levels of replication that are ∼1000-fold higher than those of WT mice, indicating that the adaptive immune system plays a significant role in reducing viral replication to enable the establishment of the chronic-phase set-point. Determining why this relative degree of adaptive immune control takes so long (∼100 days) to develop, and why further control beyond the set-point is not possible in most WT animals, will likely require the development of PgV-specific immunologic reagents (e.g., antibodies, tetramers). However, our studies in IFN-deficient mice provide some initial clues. Specifically, elevated type-I IFN signaling is known to interfere with effective T cell response in chronic viral infections (*15, 23*), and the high rate of maPgV clearance in *IFNAR*^-/-^ mice suggests that type-I IFN-induced T cell dysfunction may also be playing a role in PgV persistence. This could explain the higher prevalence of HPgV infection in people with cancer and HIV infection, as chronic elevation of type-I IFN signaling is observed in both of these conditions (*15*). However, whether PgV is causing elevated type-I IFN signaling or merely benefiting from its presence remains unknown and, in any event, does not explain the positive impact of PgV infection on HIV disease progression.

The PD-1/PD-L1 axis is another important host-determinant of persistence for many viral infections, but our data indicates that PD-1 signaling contributes minimally to PgV persistence. This is perhaps reassuring given the prevalence of HPgV and the rising frequency of PD-1 “checkpoint inhibitor” therapies for treating cancer. But more broadly, this finding implies that PgVs employ a highly novel mechanism of immune evasion that is independent of the PD-1/PD-L1 tolerance axis that is a hallmark of many chronic viral infections.

## METHODS

### Historical perspective

Initial studies adapting RPgV to infect mice were conducted at Washington University in St. Louis in 2018-2019 (Figure 1A,B); however, these were interrupted by the COVID-19 pandemic. All subsequent studies were performed at the University of Wisconsin–Madison between 2021 and 2024.

### Ethics statement

All experiments were conducted by trained personnel and in compliance with Washington University in St. Louis and the University of Wisconsin–Madison policies and procedures.

#### Mice

All mice were obtained from Jackson Laboratories and (Bar Harbor, ME) and bred in the Mouse Breeding Core at UW–Madison.

| Abbr. | Technical name | Jax ID |
| --- | --- | --- |
| WT | c57BL6/J | 000664 |
| STAT1<sup>-/-</sup> | B6.129S(Cg)-Stat1<sup>tm1Dlv</sup>/J | 012606 |
| IFNAR<sup>-/-</sup> | B6(Cg)-Ifnar1<sup>tm1.2Ees</sup>/J | 028288 |
| IFNGR<sup>-/-</sup> | B6.129S7-Ifngr1<sup>tm1Agt</sup>/J | 003288 |
| IFNAGR<sup>-/-</sup> | B6.Cg-Ifngr1<sup>tm1Agt</sup>Ifnar1<sup>tm1.2Ees</sup>/J | 029098 |
| PD1<sup>-/-</sup> | B6.Cg-Pdcd1<sup>tm1.1Shr</sup>/J | 028276 |

### 5′ RACE

The SMARTer® RACE 5’/3’ kit (Takara Bio, USA) was used to generate cDNA according to the manufacturer’s instructions using a maPgV-specific reverse primer (TGCGAGAGCCGTCAGCCACA). PCR amplification was then performed with the kit’s universal primer mix and a second maPgV-specific reverse primer (CCGTAGCAGGCGGGTCAGCA).

Finally, nested PCR was performed using the SMARTer® RACE 5’/3’ kit’s short universal primer and a third maPgV-specific reverse primer (CGCGCAAGCCCTTCTGGATA), yielding a product of ∼600bp. This PCR product was submitted for Sanger sequencing (UW Biotechnology Center) using a fourth maPgV-specific reverse primer (TCCGGCGTGGTTGTTGTGTTT), yielding high-quality chromatograms that clearly identified the “template-switching oligo” indicative of the 5′ end of the maPgV genome.

### 3′ RACE

Extracted maPgV RNA was subjected to poly(U) tailing using a poly-U polymerase (NEB, Ipswitch, MA) followed by cDNA synthesis with Superscript IV RT (Invitrogen, Waltham, MA) and a poly-A primer (GAATCGAGCACCAGTTACGCATGCCGAAAAAAAAAAAAAAAAAAAAAMN). PCR amplification was then performed using the Platinum SuperFi II DNA polymerase and a maPgV-specific forward primer (GGGGTTGGCCAGCCGATTGT) with a reverse primer complementary to the sequence added during cDNA synthesis (GAATCGAGCACCAGTTACG). This product was then subjected to nested PCR using a second maPgV-specific forward primer (CCGGCTCGGTTCAGCCATCC) and a third reverse primer (GAGCACCAGTTACGCATGCC). The nested PCR product was then subjected to Oxford Nanopore sequencing (Plasmidsaurus, Eugene, OR) which revealed homology to the known maPgV 3′ genome extending into a poly-T tract indicative of the distal 3′ end of the genome initially labeled by the poly-U polymerase.

### RNA structure analysis (Hyde)

### Mature peptide prediction (Grove)

### LCMV

A stock of passage 5 Lymphocytic choriomeningitis virus Clone 13 was provided by Arthur Kim (The Scripps Research Institute). *PD1*^-/-^ mice aged 7-11 weeks were inoculated intravenously with 2×10^6^ focus forming units diluted in sterile PBS.

#### RNA extraction

RNA was extracted from 20μL of serum using the KingFisher Flex (ThermoFisher, Waltham, MA) with MagMax reagents. Carrier RNA was omitted for cellular samples, samples destined for RACE, and samples destined for sequence-independent single-primer amplification (SISPA) sequencing, but carrier RNA was included for serum samples destined for RT-qPCR or Primal sequencing.

#### RT-qPCR

Extracted RNA was subjected to RT-qPCR using the TaqMan RNA-to-CT 1-Step Kit (ThermoFisher, Waltham, MA) in a 20μL reaction with 0.5μM of primers and 0.1μM of probe labeled with FAM and ZEN/Iowa Black quenchers (IDT, Coralville, IA). Primer/probe sets for RPgV/maPgV were as follows: Forward: ATCACGGGTAAGCTGGTTTG; Reverse: GGAAACCAAGCAGAGTGAGC; Probe: CGGACACTTCCCAGTCTGT. Primer/probe sets for HPgV were as follows: Forward: GGCGACCGGCCAAAA; Reverse: CTTAAGACCCACCTATAGTGGCTACC; Probe: TGACCGGGATTTACGACCTACCAACCCT. Thermocycling was performed on a Quantstudio 6 Pro (Applied Biosystems, Waltham, MA) with a 96-well block (0.2mL) under the following conditions: 48°C for 15 min followed by 95°C for 10 min, then 50 cycles of 95°C for 15 sec followed by 60°C for 1 min. A RNA standard was made by cloning a fragment of the maPgV genome sequence into the pJET1.2/blunt vector (Invitrogen, Waltham, MA). After linearization of the construct, transcription was performed in vitro for 6 h with the MEGAscript T7 transcription kit (Invitrogen, Waltham, MA), followed by purification using the MEGAclear transcription cleanup kit (Invitrogen, Waltham, MA), quantification, and dilution to a concentration of 1×1010 transcript copies per μL. Ten-fold dilutions of this transcript were used as a standard curve, which was linear over 8 orders of magnitude and sensitive down to 10 copies of RNA transcript per reaction.

#### Unbiased deep sequencing

cDNA was generated from extracted RNA using a revised SISPA approach. First, 30 μL of extracted total nucleic acids were treated with TURBO DNase (Thermo Fisher Scientific) and concentrated to 10 μL with an RNA Clean & Concentrator-5 kit (Zymo Research, Irvine, CA, USA). Next, 1 μL of Primer A (40 pmol/µL; 5′-GTTTCCCACTGGAGGATA-(N9)-3′) was added to 4 μL of concentrated viral RNA and heated in a thermocycler at 65°C for 5 min and cooled at 4°C for 5 min. Reverse transcription was performed by adding 5 µL of Superscript IV (SSIV) reverse transcription master mix containing 1 µL of deoxyribonucleotide triphosphate (dNTP; 10 mM), 0.5 µL of dithiothreitol (DTT; 0.1 M), 1 µL of PCR water, 2 µL of 5X RT buffer, and 0.5 µL of SSIV RT to the reaction mix. The mix was incubated in a thermocycler at 42°C for 10 min. Second-strand cDNA synthesis was performed by adding 5 µL of Sequenase reaction mix (3.85 µL of PCR water, 1 µL of 5X Sequenase reaction buffer, and 0.15 µL of Sequence enzyme) to the reaction mix and incubating at 37°C for 8 min. After incubation, 0.45 µL of the Sequenase dilution buffer and 0.15 µL of Sequenase were added to the reaction mix and incubated at 37°C for 8 min. To amplify the cDNA, 5 µL of the cDNA was added to 45 µL of the Primer B reaction mix containing 0.5 µL of AccuTaq LA DNA polymerase, 5 µL of AccuTaq LA 10x buffer, 1 µL of Primer B (100 pmol/µL; 5′-GTT TCC CAC TGG AGG ATA -3′), 2.5 µL of dNTP (10 mM), 1 µL of dimethyl sulfoxide (DMSO), and 35 µL of PCR water. The cDNA was amplified using the following thermocycler conditions: 98°C for 30 s, 30 cycles (94°C for 15 s, 50°C for 20 s, and 68°C for 2 min), and 68°C for 10 min. After the incubation, the amplified PCR product was purified using AMPure XP beads (Beckman Coulter, Brea, CA, USA) at a 1:1 concentration and eluted in 50 µL of PCR water. The purified PCR products were quantified with the Qubit dsDNA high-sensitivity kit (Invitrogen, Waltham, MA, USA). SISPA-prepared cDNA material was submitted to the University of Wisconsin–Madison Biotechnology Center. Samples were prepared according to the QIAGEN FX DNA Library Preparation Kit (QIAGEN, Germantown, MD, USA). The quality and quantity of the finished libraries were assessed using a Tapestation (Agilent, Santa Clara, CA, USA) and a Qubit dsDNA HS Assay Kit, respectively. Paired-end 150-bp sequencing was performed using the NovaSeq6000 system (Illumina, San Diego, CA, USA).

#### Primal sequencing

Primers were designed to generate overlapping amplicons of ∼250bp spanning the entire maPgV genome using PrimalScheme in high-GC mode (*24*). Upon optimization, primer pools were spiked with additional primers, designed manually in Primer3_v.0.4.0 (*25, 26*), to improve read depth in areas with reproducibly poor coverage. Illumina TruSeq adaptors were added to maPgV-specific primers and primers were ordered in “even” and “odd” oPools based on sequential amplicon number so as to minimize interference between overlapping amplicons during PCR amplification (IDT, Coralville, IA). RT-PCR was performed on extracted maPgV RNA according to recommendations from PrimalScheme using the SuperScript IV One-Step RT-PCR System (Invitrogen, Waltham, MA). Following cleanup with Ampure XP beads (Beckman Coulter, Brea, CA), “even” and “odd” amplicon pools were combined for each sample and subjected to index PCR (UW Biotechnology Center). Paired-end 150-bp sequencing was performed using the NovaSeq6000 system (Illumina, San Diego, CA, USA).

#### Sequence analysis

Illumina sequencing data was imported into Geneious Prime (version 2022.2.2) (Biomatters, Auckland, New Zealand) as paired reads, then merged using the BBMerge tool Version 38.84. Merged reads were then mapped to the “consensus RPgV” sequence––this reference was used for all mappings and is comprised of the RPgV consensus sequence with the first 156 nucleotides originating from maPgV 5′ RACE and the last 151 nucleotides (nt 10637–10941 in the genome) originating from maPgV 3 ′ RACE. Read mapping was performed using Bowtie2 (v2.4.5) (*27, 28*), with 20bp (for sequences generated via unbiased/SISPA) or 30bp (for sequences generated via PrimalScheme) trimmed from both ends prior to mapping to remove primers. Single nucleotide variants were then characterized using the built-in “find variations/SNPs” function in Geneious, using a minimum variant frequency of 0.05 and a minimum coverage of 100.

#### Data availability

Raw datasets used to make each figure will be uploaded to Dryad and associated with Adam Bailey’s ORCiD: 0000-0002-6560-9680.

## ACKNOWLEDGEMENTS

We are extremely grateful to Dr. Michael Diamond (Washington University in St. Louis) whose lab provided the mice and resources that allowed for the initial adaptation of RPgV to maPgV. Amit Kapoor (Ohio State University) provided the original RPgV+ sample. We also thank the University of Wisconsin School of Medicine and Public Health Biomedical Research Model Services shared resources for the use of its facilities and their Research Services team for expertise including mouse breeding. The authors utilized the University of Wisconsin–Madison Biotechnology Center for deep sequencing and mass spectrometry.

---

**Figure S1.**
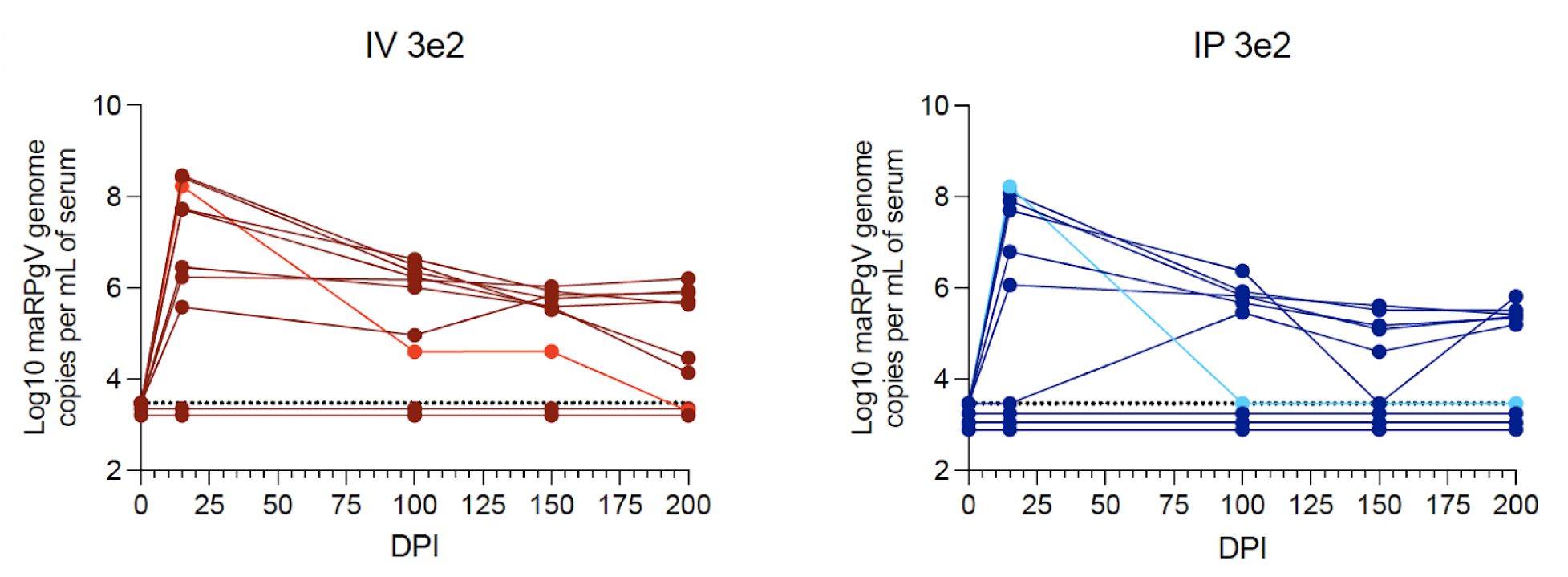
Repeat of low-dose inoculation studies.

**Figure S2.**
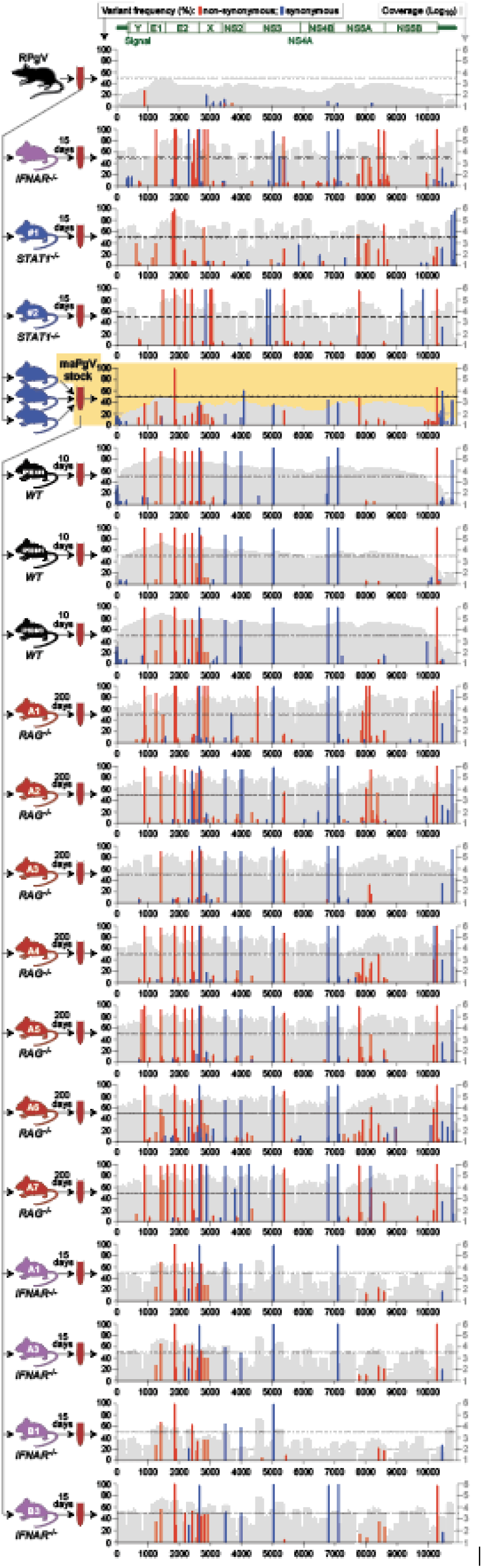
Expanded sequencing cohort.

